# Antibody full-length light chain reshapes variable domains aggregation without altering their dominance in amyloid fibrils

**DOI:** 10.64898/2026.11.16.766878

**Authors:** Basudha Patel, Sharvari Palkar, Amit Kumawat, Annesha Ghosh, Shivansh Singh, Twinkle Aggarwal, Swasti Pradhan, Saara Syam, Sandhya Gokula Nathan, Jogender Singh, Sarita Puri

## Abstract

AL amyloidosis is a fatal systemic amyloid disease in which antibody light chains (LCs), including full-length (FL) and their proteolytically fragmented variable (VL) and constant (CL) domains, aggregate within a complex extracellular matrix (ECM) across multiple tissues. However, how these co-existing species shape aggregation kinetics, fibril morphology, stability and toxicity remain unexplored. In this work, using a patient-derived LC (AL55), we show that FL transiently interacts with VL, slowing its aggregation nearly four-fold and diverting it into protease-sensitive, seeding-incompetent protofibrillar aggregates rather than mature amyloid assemblies. VL alone or in the presence of inert CL instead matures into toxic, protease-resistant, seeding-competent helical fibrils. Owing to their greater protease resistance and seeding competence, VL-dominated amyloid assemblies persist and contribute to their enrichment within tissues. The FL-mediated redirection is generic across multiple amyloidogenic VLs. These results reveal that LC aggregation, like that of Aβ and Tau, is governed by interplay between full-length and fragmented species, rather than VL domain aggregation in isolation. Overall, this work identifies a molecular basis for the selective enrichment of VL fibrils in patients and provides a therapeutic rationale for stabilizing VL domains to prevent their pathological aggregation.

**Significance statement:** AL amyloidosis deposits contain FL LCs and their proteolyzed VL and CL-domain fragments, yet how the presence of FL and CL domain influences aggregation kinetics, fibrillar morphology, and toxicity of amyloidogenic VL domain remains unexplored. This study resolves this gap by showing that the FL LC transiently engages with VL, slowing its aggregation and redirecting it into fragile, non-toxic, seeding-incompetent protofibrils, whereas free VL matures into rigid, toxic, seeding-competent fibrils that persist and dominate patient-derived amyloid deposits.

## INTRODUCTION

AL amyloidosis, one of the most prevalent systemic amyloidoses, arises from the misfolding, aggregation, and extracellular deposition of monoclonal immunoglobulin light chains (LCs) secreted by clonally expanded plasma cells (Merlini et al., 2018; Blancas-Mejía et al., 2018; Quock et al., 2018). Amyloid accumulation in vital organs such as the heart and kidneys disrupts tissue architecture and function, impairing cardiac contractility and glomerular filtration, and leading to rapid organ failure and high mortality (Li et al., 2019). Defective plasma cells secrete native full-length (FL) LCs mostly as homodimers, in which each monomer comprises a variable (VL) and a constant (CL) domain (Bourne et al., 2002; Oberti et al., 2017). Aggregation occurs within the extracellular matrix (ECM) of individual organs, where proteolysis, occurring either before or after aggregation, contributes to the formation of VL domain-dominated amyloid assemblies (Radamaker et al., 2021; Swuec et al., 2019; Radamaker et al., 2019; Radamaker et al., 2021; Puri et al., 2023; Yu et al., 2025; Yao et al., 2026; Schulte et al., 2024; Hicks et al., 2026; Bassett et al., 2026). These observations are consistent with in vitro aggregation studies and aggregation inside an AL mouse model showing that FL LCs and truncated CL domains do not aggregate unless conditions are drastically altered, for example, by acidic pH or the addition of detergents such as sodium dodecyl sulphate (SDS) or a denaturant such as urea or addition of pre-formed VL aggregates as seed, whereas isolated VL domains readily aggregate under native conditions (Andrich et al., 2107; Rennella et al., 2019; Puri et al., 2025c; Lavatelli et al., 2024; Martinez-Rivas et al., 2025).

In contrast, crude organ extracts from AL patients form heterogeneous deposits containing FL LCs together with proteolytically generated VL and CL fragments of varying lengths, serum amyloid P protein, and ECM components such as collagen (Buxbaum et al., 2022; Gottwald et al., 2021; Lavatelli et al., 2020; Lavatelli et al., 2008; Marchese et al., 2026; Schulte et al., 2024). These observations are also consistent with findings from the recently developed mouse model by the Sirac group, in which fibril extracts from the heart of the AL mouse consist of FL LCs together with fragmented domains of varied lengths (Martinez-Rivas et al., 2025). Moreover, no aggregation and deposition was observed upon sole over-expression of a full-length amyloidogenic light chain until supplemented with the VL derived in-vitro preformed seeds (Martinez-Rivas et al., 2025), suggesting proteolysis is an important step to release aggregation competent VL domain which can sequester full-length LC and induce its aggregation. Similar effects may arise either from non-specific sequestration of ECM components by hydrophobic or charged residues exposed on the surface of amyloid assemblies or from specific interactions that could modulate aggregation pathways, fibrillar structure, and composition.

Mass spectrometry and high-resolution cryo-electron microscopy (cryo-EM) structures obtained from ex vivo purified fibrils have shown that VL domains predominantly form the amyloid cores (Radamaker et al., 2021; Swuec et al., 2019; Radamaker et al., 2019; Radamaker et al., 2021; Puri et al., 2023; Yu et al., 2025; Yao et al., 2026), with only rare or limited extensions into CL regions (Schulte et al., 2024). This disconnect between the biochemical heterogeneity of patient deposits and the structural homogeneity observed in cryo-EM structures has led to reduced emphasis on investigating how FL LCs and their truncated fragments modulate LC aggregation. Instead, studies have largely focused on highly aggregation-prone VL domains in isolation (Meunier-Carmenate et al., 2023; Pradhan et al., 2023; Wong et al., 2024; Puri et al., 2025c; Kazman et al., 2021), a direction that may overlook the complex aggregation in vivo.

Here, we address this gap by studying VL domain aggregation in presence of FL and truncated CL domain using a patient-derived amyloidogenic light chain (AL55). AL55 belongs to the λ6 germline, the most common germline among AL patients (Puri et al., 2025a; Lavatelli et al., 2020; Puri et al., 2025b; Puri et al., 2025c; Paissoni and Puri et al., 2025). This protein forms fibrillar deposits across multiple organs, and the associated patient died of cardiac failure (Swuec et al., 2019; Puri et al., 2023). CryoEM studies on the purified fibrils showed that VL domain consisting of residues 1 to 111 form the identical fibrillar core in multiple organs like heart and kidneys (Swuec et al., 2019; Puri et al., 2023). Moreover, previous work from our group, as well as Lavatelli et al., has shown that the FL and isolated CL domain of AL55 are resistant to aggregation under in vitro conditions, whereas the VL domain forms single-protofilament containing helical assembly (Puri et al., 2025b; Lavatelli et al., 2024). Crude extracts from the patient, by contrast, showed that amyloid clumps comprise a mixture of FL AL55 and its fragmented VL and CL domains of varying lengths (Swuec et al., 2019; Lavatelli et al., 2020). Under in vitro conditions, AL55 VL interacts with extracellular chaperones apolipoprotein E and clusterin, which modulate its aggregation kinetics, fibrillar morphology, and stability (Marchese et al., 2026).

AL55 therefore provides a well-characterized model system to study modulation of VL domain aggregation and its fibrillar outcomes in presence of FL and CL domain and to simply the system, we used FL, VL, and CL domain for current work and have not included multiple fragment lengths among VL and CL domains. By integrating aggregation kinetics assays, fibrillar composition and structural comparisons, proteinase K (PK) digestion, cross-seeding aggregation assays, cryo-EM, and toxicity assays, we show that the isolated VL domain, alone or in the presence of inert CL, aggregates faster and forms long, stable, helical, seeding-competent fibrils. These fibrils exert significant toxicity by reducing pharyngeal pumping in *C. elegans*. In contrast, aggregation of VL in presence of FL AL55 reduces its aggregation rates by nearly four-fold, and redirects its fibrillar structure toward proteolytically labile, seeding-incompetent, non-toxic protofibrils.

Together, these findings provide a mechanistic framework that reconciles the heterogeneity of LC species observed in patient tissues with the selective visualization of VL-derived fibril structures in cryo-EM studies. Our results establish that LC aggregation is a complex process where presence of FL protein and its truncated domains in a confined space modulate aggregation rates, fibrillar stability, proteolysis sensitivity, and seeding competence significantly. Together, these properties govern which assemblies persist through extraction and dominate amyloid deposits, positioning VL domain amyloid assemblies as the stable, extractable forms as found in AL patients.

## RESULTS

### Model proteins

FL AL55 is the first reported open homodimer structure among all known LC structures, in which two VL domains do not interact with each other (Puri et al., 2025b). Sequentially, each monomer consists of a VL domain (residues 1-111) and a CL domain (residues 112-217) (Figure S1A). The crude AL55 fibrillar extract from the patient is a mixture of FL and truncated VL and CL domains of varying lengths, as shown by Lavatelli et al. (Figure S1B), while VL from residues 1-111 form the fibrillar core (Lavatelli et al., 2020; Swuec et al., 2019; Puri et al., 2023; Lavatelli et al., 2008). Moreover, under in vitro conditions, only the VL domain undergoes aggregation, while FL and CL are resistant to aggregation (Figures S1C, S2) (Lavatelli et al., 2024; Puri et al., 2025b). Therefore, we used recombinantly purified FL, VL (residues 1-111), and truncated CL domain as a model to study impact of FL and CL on aggregation of VL domain. All three model proteins were purified from the host *E. coli* (Figure S3) and correctly folded pure proteins were used to aggregation assays (Figure S4).

### Aggregation kinetics assays show FL slows VL aggregation

Ex vivo aggregates containing FL and truncated domains of varied lengths may result from non-specific interactions between native or misfolded forms of these proteins, or from native proteins decorating mature fibrils (Swuec et al., 2019; Lavatelli et al., 2020; Schulte et al., 2024;). However, it is not known how truncation and the presence of truncated fragments in a confined space modulate the aggregation process and its outcome. Therefore, to map the effect of FL and CL domains on aggregation kinetics of the aggregation-prone VL domain, we performed aggregation kinetics assays at 1:1 and 1:2 molar ratios of VL:FL and VL:CL proteins. We found that FL reduced the aggregation rate by approximately four-fold, from 0.063±0.005 min^-1^ to 0.017±0.0008 min^-1^, in a concentration-dependent manner. This was accompanied by a linear increase in lag time from 35±5 to over 300 minutes (Figures 1A-B). The delayed lag phase indicated the conformational changes occurring in VL domain to reach to aggregation competent forms and their accumulation in large amounts to initiate the aggregation (nucleation) are delayed probably due to FL-VL interactions (Arosio et al., 2015). The reduced ThT fluorescence intensity at the end of the aggregation reaction suggested that either the amount or the morphology of the fibrils formed was affected. Supernatant and pellet fractionation of the aggregation products confirmed a concentration-dependent reduction in aggregated VL protein, with a visible reduction in higher molecular weight (MW) populations (Figure 1C). Moreover, the presence of the faint FL band suggested either its sequestration or co-aggregation with VL.

**FIGURE 1.**
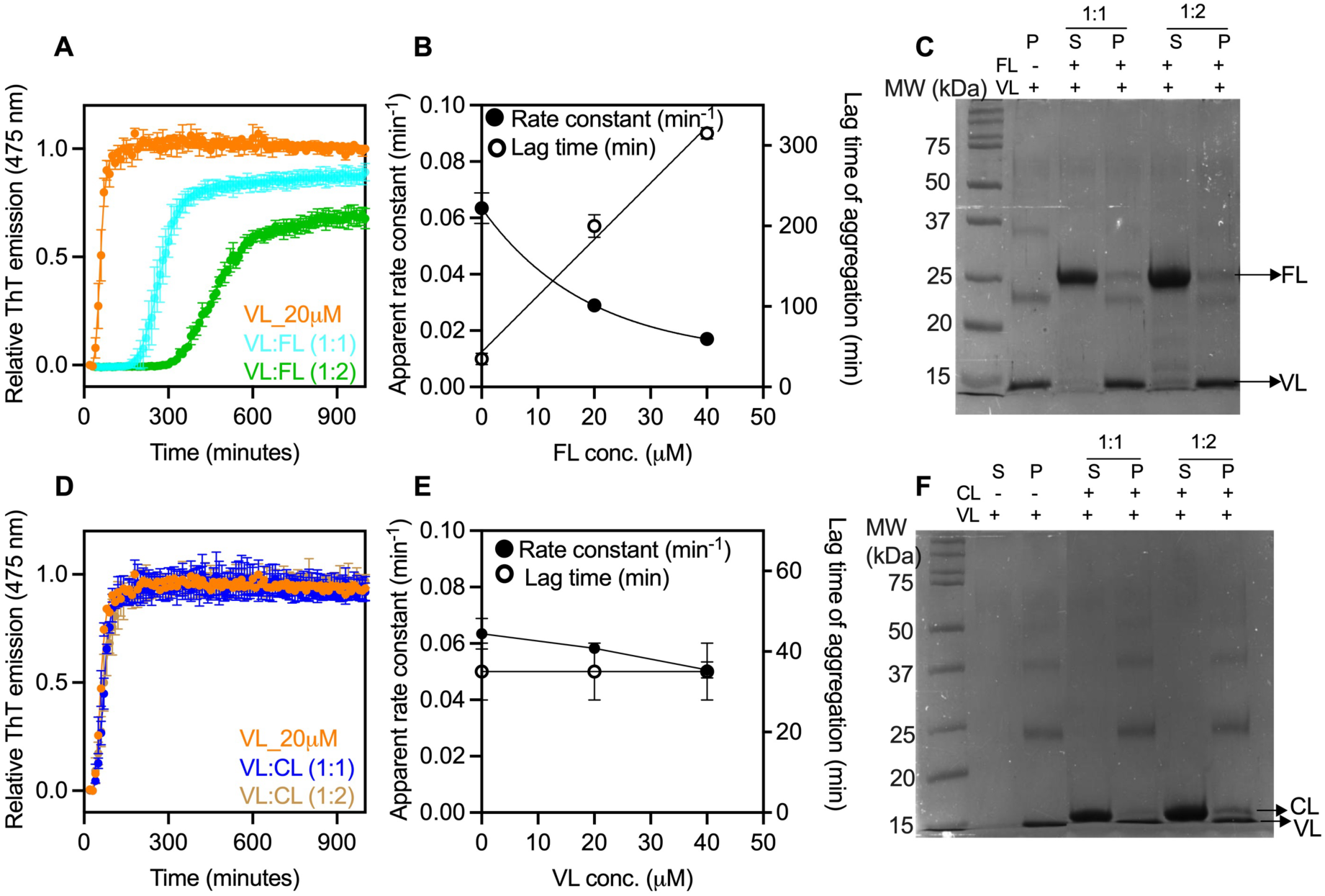
**Aggregation kinetics assay of VL in presence of FL and CL domain.**(A) Aggregation kinetics assay of VL in the presence of FL in a molar ratio of 1:1 and 1:2 (VL:FL). The error bars in each aggregation profile represent the heterogeneity in the ThT fluorescence emission intensity among different replicates. (B) Correlation plot between apparent aggregation rate constants and lag time with respective FL concentrations. (C) Supernatant and pellet fractionation of aggregated samples containing VL-alone, VL:FL (1:1), and VL:FL (1:2) molar ratios. (D) Aggregation kinetics of VL in the presence of CL in a molar ratio of 1:1 and 1:2 (VL:CL). (E) Correlation plot between aggregation rate constants and lag time with the respective CL concentrations. (F) Supernatant and pellet fractionation of aggregated samples containing VL-alone, VL:CL (1:1), and VL:CL (1:2) molar ratios. Although both VL and CL have similar molecular weights of 11-12 kDa, CL runs slightly slower in the SDS-PAGE, a common feature of CL domains due to high amount of lysine and arginine in their sequence. CL has 9 lysine residues and VL has only three. The isoelectric points (pI) of VL and CL proteins are 4.87 and 6.91, respectively.

In contrast, the presence of the CL or bovine serum albumin (BSA) did not affect the aggregation profile of VL in terms of aggregation rates and lag time (Figures 1D-E, Figure S5). Supernatant and pellet fractionation showed similar amounts of aggregated VL with and without the CL, and most of the CL was present in the supernatant fraction (Figures 1F). Together, these experiments show that FL significantly modulates VL aggregation, whereas CL has no effect.

### FL restricts VL aggregates to protofibril forms

Next, we compare the fibrillar architectures using transmission electron microscopy (TEM) to understand how different aggregation conditions affect fibril assembly. Isolated VL and VL aggregated in presence of CL resulted in long fibrils within 24 h (Figure S6). These fibrils further matured into polymorphic helical fibrils with widths varying between 8–10 nm and crossover lengths of 50-100 nm upon longer incubation for two weeks (Figures 2A-C). These features were consistent with previously reported in vitro-formed fibrils of AL55 VL (Puri et al., 2025b). Moreover, similar fibril morphology was formed under VL-alone and in presence of CL, suggesting that the CL domain does not influence fibril architecture, consistent with its negligible effect on aggregation kinetics (Figure 2C). In contrast, the presence of FL markedly altered fibril morphology, favoring the formation of amorphous aggregates at short aggregation time (Figure S6) which mature to protofibrillar assemblies with a thickness of 4-5 nm rather than mature fibrils of 8-12 nm typical for LC fibrils (Figure 2B). We also collected the cryo-EM data of protofibrils to obtain further information about their structural organization and heterogeneity (Figure 2D and Figure S7). 2D class averages revealed polymorphic protofibrils exhibiting straight (29.6%), twisted (16.5%), bent (48.8%) and additional poorly resolved species composed of β-sheet structures. The twist length in the twisted polymorph was about 500 Å (Figures 2E-F). The distance between individual β-strands was estimated to be 4.75 Å from the helical layer-line plot obtained using RELION, a typical feature of amyloid assemblies (Figure 2F and Figure S7). Owing to the lack of helicity, we were unable to solve high-resolution 3D structures of the protofibrils. Collectively, these findings demonstrated that FL remodels VL aggregation significantly by promoting the formation of structurally distinct protofibril-rich aggregates in LCs.

**FIGURE 2.**
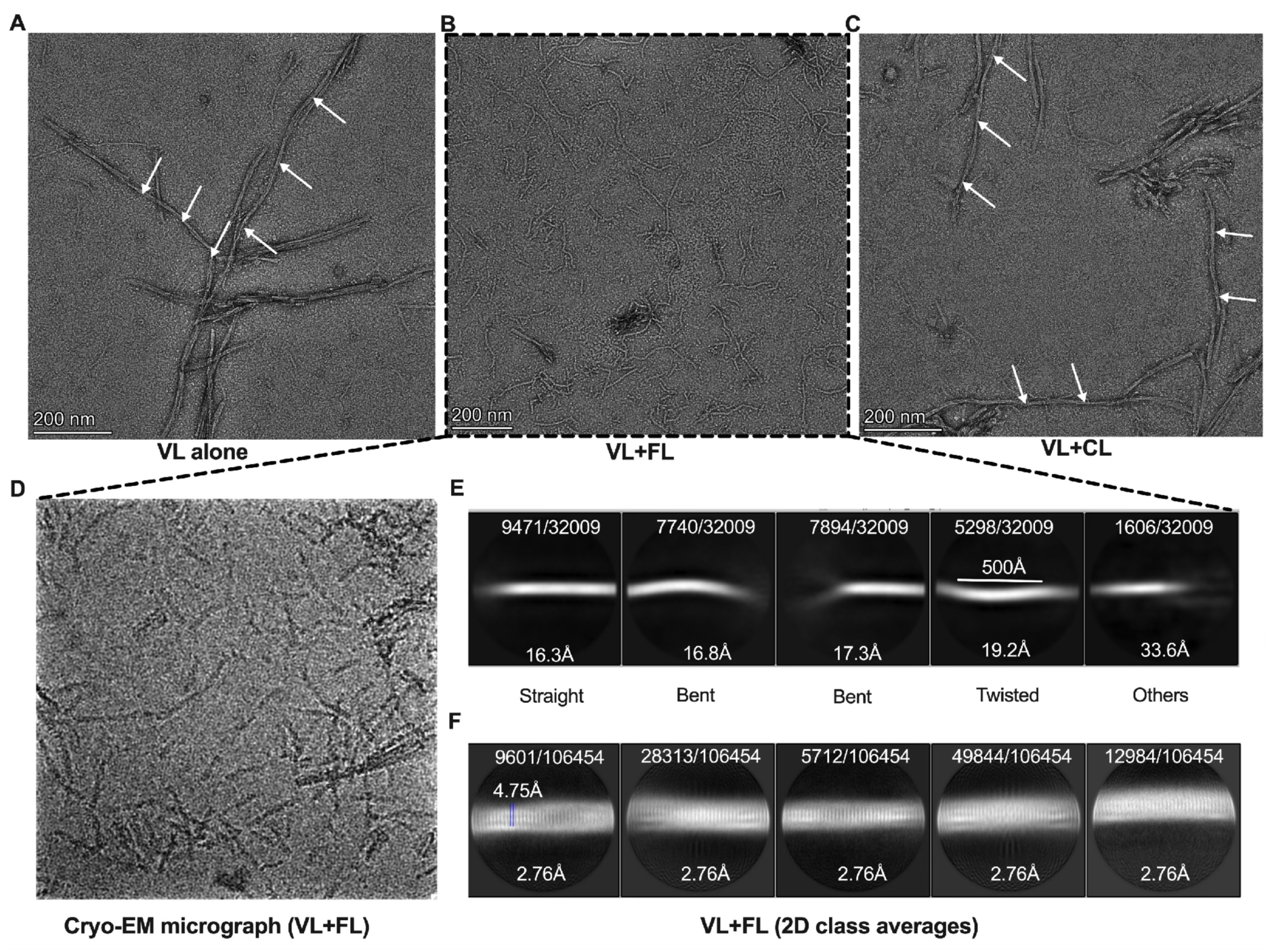
Fibrillar architecture comparison under different aggregation conditions. (A) Negative stain TEM data of VL-alone fibrils after 15 days of aggregation reaction at a magnification of 46 k. (B) VL+FL aggregation reaction protofibrils. (C) VL+CL aggregation reaction fibrils. The white arrows represent the helical pitch distance of different fibrils. (D) Representative cryo-EM micrograph of protofibrils at a magnification of 120 k. (E) Large 2D class averages obtained from the cryo-EM data of VL+FL protofibrils with a box size of 768 pixels. (F) Small 2D class averages obtained from small box size (280 pixels) extracted particles. We can clearly visualize the β-strand separation at a distance of 4.75 Å. The overall resolution of all classes is 2.76 Å.

### VL is the major fibrillar component in all aggregation reactions

To assess the presence of FL and CL proteins within VL aggregates over time, we first analyzed pellet fractions collected after 24 h and 15 days of aggregation using SDS-PAGE. SDS–PAGE analysis of the 24 h samples revealed that the VL+FL aggregation sample was a mixture of VL and FL proteins, whereas the VL+CL aggregation sample was predominantly composed of VL at this early time point (Figures S8A-B). The presence of a small amount of FL along with VL in the pellet fraction might be a result of its non-specific sequestration. This was confirmed by the SDS-PAGE gel run after 15 days of the aggregation reaction. Both VL+FL and VL+CL aggregation samples contained exclusively VL in their pellet fractions, while FL and CL proteins were detected in the supernatant and/or appeared degraded (Figure S8C). This was further confirmed by the size-exclusion chromatography (SEC) profile of the supernatant fraction obtained from VL+FL aggregation reaction after 24 h (Figure 3A). The high-molecular-weight peak eluting around 9.0 ml corresponded solely to VL soluble oligomers, as confirmed by SDS-PAGE, indicating that FL did not participate in aggregation. Additionally, the major peak corresponded to native FL, confirming that most of the FL was present in the supernatant (Figures 3A-B, Figure S8A). Together, these results indicated that FL engaged in transient interactions with native VL during the early stages of aggregation, modulating VL aggregation rates and fibrillar morphology, but did not participate in its oligomerization and aggregation. Moreover, the lack of detectable interactions between VL and CL, in either the native or fibrillar state, permitted unperturbed VL aggregation (Figure S8B), resulting in fibrillar assemblies indistinguishable from those formed by VL-alone.

**FIGURE 3.**
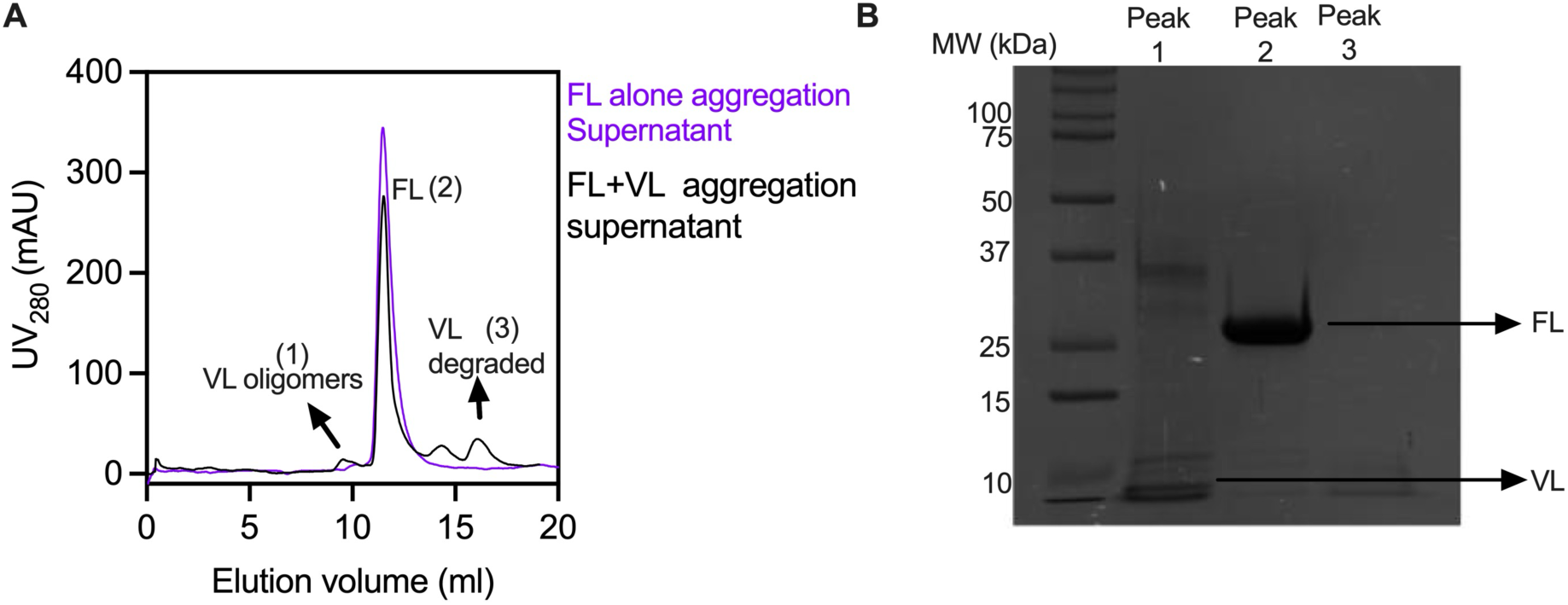
**Composition of fibrils under different aggregation assays**. (A) SEC profile of supernatant collected after 24 h of VL+FL mixed-aggregation reaction (black) and FL alone (magenta). A small peak at elution volume 9 ml is transiently populated VL soluble oligomers, as confirmed by SDS-PAGE gel (Panel B). (B) SDS-PAGE analysis of three collected peaks named as 1, 2, and 3; where peak 1 contains oligomeric VL, peak 2 is soluble FL, and peak 3 is a small amount of degraded VL.

### VL protofibrils are sensitive to proteolysis and incompatible with cross-seeding VL aggregation

Further, to compare stability and cross-seeding competence of fibrils formed under different aggregation reactions, we performed a limited PK digestion assay and cross-seeding aggregation kinetics assays. Without PK digestion, VL fibrils formed in ThT-based aggregation assays appeared as multiple oligomeric species on SDS– PAGE. Such behavior is well documented for AL55 VL and other amyloidogenic VL domain fibrils, which are resistant to ionic detergents such as SDS, the denaturant urea, and heat treatment, and typically display multiple bands on SDS–PAGE (Pradhan et al., 2023; Puri et al., 2025c). Upon PK treatment, higher-molecular-weight species were proteolyzed, while lower-molecular-weight species persisted. A similar pattern was observed for fibrils from the VL+CL aggregation reaction, indicating that VL fibrils formed alone or in the presence of CL exhibit comparable stability and PK sensitivity. In contrast, aggregates from the VL+FL aggregation reaction were more sensitive to PK digestion, showing digestion of up to 60-70% at both 1:200 and 1:500 (PK:protein) ratios (Figure 4A; Figure S9). The higher proteolysis rate of these unstable species might contribute to their faster clearance by the extracellular proteolytic machinery, compared with the less sensitive VL-alone and VL+CL aggregation reaction fibrils, which accumulate and are present in large amounts in patient extracts (Swuec et al., 2019; Lavatelli et al., 2020).

**FIGURE 4.**
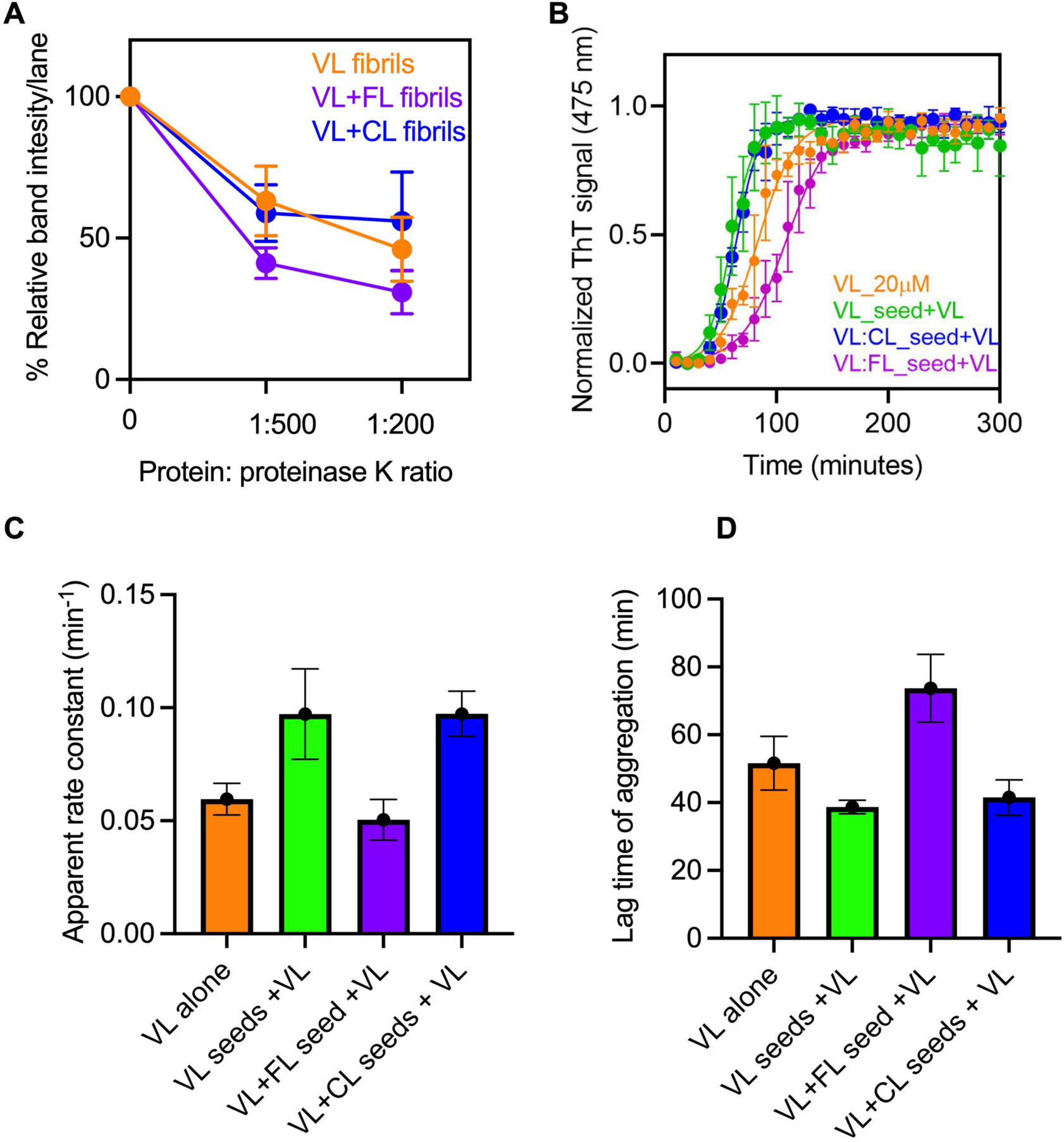
Comparison of fibril stability and cross-seeding compatibility. (A) Correlation plot of relative band intensities per lane vs PK:protein ratio obtained from three independent proteinase K digestion assays performed on the aggregated samples obtained from three different days reactions. (B) Cross-seeding aggregation kinetics of isolated VL domain (orange) in the presence of VL only (bright green), VL+FL (magenta), and VL+CL mixed aggregation seeds (blue). The experiment was performed in three technical replicates. (C-D) Histogram showing apparent aggregation rate constants and lag time of aggregation for VL-alone, VL seeded with VL fibrils, VL cross-seeded with VL+FL protofibrils, and VL cross-seeded with VL+CL fibrils. The error bars are standard error measurements from three technical replicates. The colour coding is the same as panel B.

Moreover, in patients, heterogeneous light chain assemblies may initially form; however, only species capable of efficient seeding are likely to persist and expand, whereas poorly seeded intermediates are cleared or remodeled (Morfino et al., 2022). To test this hypothesis, we performed cross-seeding aggregation assays using 5.0% (w/w) preformed fibrils from VL-alone, VL+FL, and VL+CL aggregation reactions as seeds for native VL aggregation. Preformed fibrils formed from VL-alone and VL+CL aggregation reactions effectively seeded VL aggregation, resulting in an approximately 1.5-fold increase in aggregation rate compared to an unseeded reaction. In contrast, VL protofibrillar seeds formed in the presence of FL led to an approximately 0.8-fold reduction in apparent rate constant, indicating poor seeding compatibility with VL monomers. This incompatibility likely reflects differences in the fibrillar core structures formed under VL-alone and VL+CL aggregation conditions relative to the VL+FL aggregation reaction, in which the protofibrillar form was predominantly populated (Figures 4B-D). Consequently, fibrils from VL-alone and VL+CL aggregation reactions can continuously recruit native VL, leading to the preferential accumulation of VL-dominated fibrillar species within the ECM. In contrast, the limited seeding efficiency and enhanced proteolytic susceptibility of VL+FL aggregated protofibrils are likely to prevent their accumulation and persistence, providing a rationale for why the VL-based amyloid fibril population is dominantly recovered from ex vivo AL patient samples used for cryo-EM analyses (Radamaker et al., 2021; Swuec et al., 2019; Radamaker et al., 2019; Radamaker et al., 2021; Puri et al., 2023; Yu et al., 2025; Yao et al., 2026; Schulte et al., 2024).

### VL domain association with FL restricts aggregation-prone region exposure

Although FL was absent in mature protofibrils, its presence significantly modulated VL domain aggregation, pointing to transient interactions between native FL and VL. Therefore, to determine whether this transient association could influence conformational dynamics, we performed molecular dynamics (MD) simulations for apo VL and the VL+FL complex (Figures S10-11). The starting structure for apo VL was extracted from the crystal structure of FL AL55 (PDB ID: 9FVK) (Puri et al., 2025b). The model for the VL+FL complex was generated using AlphaFold3 (Abramson et al., 2024), as no experimentally determined structure of the complex is currently available. The modeled structure was consistent with our previous studies on AL55 FL, which showed that aggregation-prone regions (APRs), such as residues 38-56 and 90-103, form the VL-VL interface in the trimer-of-dimers conformation (Puri et al., 2025b). Residue-wise average root-mean-square fluctuations (RMSF) calculated from the MD simulations showed that the apo VL domain exhibited higher flexibility overall, whereas these fluctuations were significantly reduced within the interface-forming APRs (residues 38-56 and residues 90-103) in the VL+FL complex (Figure 5A), consistent with persistent hydrogen bonds between selected residues at the VL-VL interaction interface (Figure 5B) (Puri et al., 2025a; Puri et al., 2025b; Puri et al., 2025c, Paissoni and Puri et al., 2025). Thus, VL binding to the FL dimer restricted exposure of APRs in the apo VL domain, thereby slowing its aggregation. In addition, this transient interaction may alter the structure of the aggregation-competent form, contributing to the significant shift in fibrillar morphology towards protofibrils rather than mature fibrils.

**FIGURE 5.**
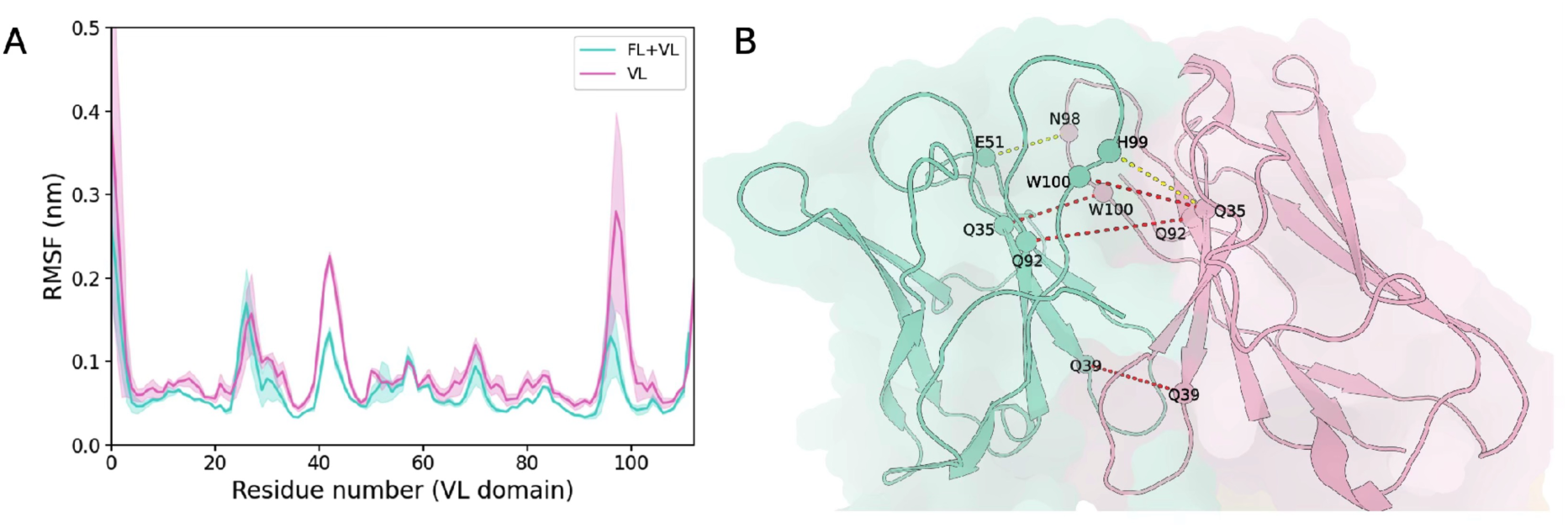
VL-domain fluctuations and interface hydrogen bonds. (A) Average RMSF plot of the VL domain in apo form and the VL+FL complex, calculated over three independent replicates. Shaded regions represent the standard deviation. (B) Representative VL-VL interaction interface in the VL+FL complex. The FL-bound VL domain is shown in teal, and the interacting VL domain is shown in pink. Hydrogen bonds are shown as dashed lines, with red and yellow indicating occupancies greater than 80% and 40%, respectively, across the three replicates.

### Rigid fibrils exert strong toxicity in *C. elegans*

*C. elegans* is a well-established model for studying cardiotoxicity of the native amyloidogenic LCs, as these affect pharyngeal pumping by causing mitochondrial stress and enhanced production of reactive oxygen species (Maritan et al., 2020; Diomede et al., 2014; Broggini et al., 2023). Here, we used the wild-type strain N2 of *C. elegans* as a model system to study the differential toxicity caused by ingested fibrils formed in VL alone, VL+FL, and VL+CL aggregation conditions. For this experiment, we fed the L4-stage larvae of *C. elegans* with 100 µg/ml of fibrils for 2 h and measured their pharyngeal pumping after 4, 24, and 48 h of growth on normal *E. coli* OP50 (CGC)-based food. Interestingly, we found that rigid fibrils formed under VL and VL+CL aggregation conditions were highly toxic, reducing pharyngeal pumping by approximately 1.5-fold, compared with a 2-fold reduction induced by the positive control, 0.1 mM H_2_O_2_ (Figures 6A-C). To further confirm the localization of ingested fibrils within the worms, we fed them with the Congo red-labelled fibrils obtained from VL-alone, VL+FL, and VL+CL aggregation reactions for 2 h at a concentration of 100 µg/ml, and observed their localization using a stereo-fluorescence microscope after 2 h. We found fewer VL alone and VL+CL reactions rigid fibrils entering the worm gut, likely owing to significantly reduced pharyngeal pumping caused by their entry (Figure 6D). In contrast, we found a large amount of VL+FL reaction protofibrils within the worm gut, with no effect on the pharyngeal pumping, suggesting that VL+FL protofibrils were not toxic to the worms. Quantitative estimation of Congo-red fluorescence intensity of ingested VL-alone, VL+FL, and VL+CL reaction fibrils is shown in Figure 6E. Together, these results demonstrate that distinct fibril morphologies exert differential uptake, mechanical stress, and toxicity in *C. elegans*.

**FIGURE 6.**
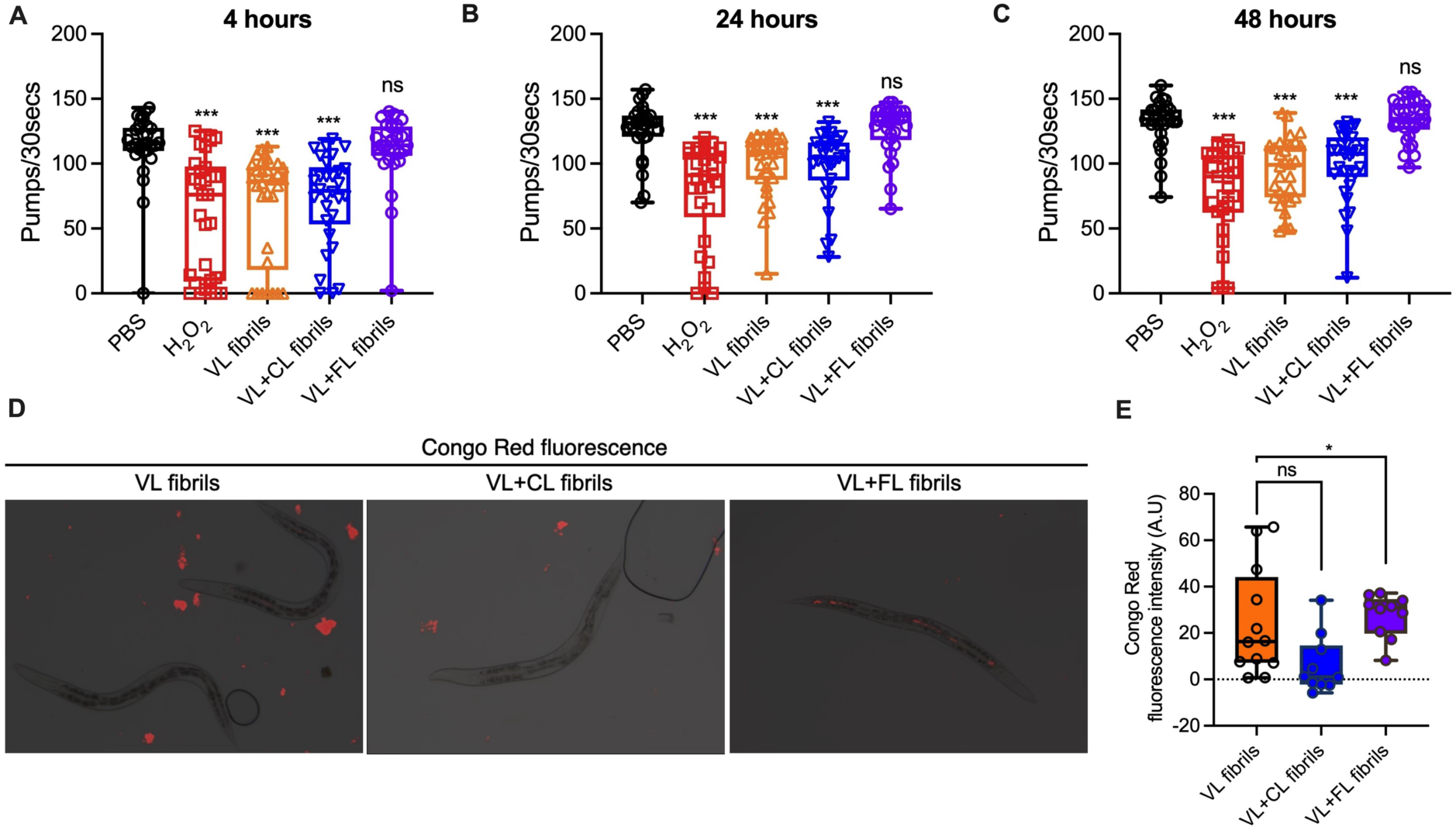
Measurements of the toxicity of different fibrillar forms. (A-C) Pharyngeal pumping measurements on 30 worms after 4, 24, and 48 h of feeding on fibrils. (D) Localization of fibrils inside *C. elegans* after 2 h of feeding. The bright red fluorescence signal corresponds to Congo-red-labelled fibrils. We observed less accumulation of VL-alone and VL+CL aggregation fibrils, but a large accumulation throughout the gut for VL+FL aggregation reaction protofibrils. (E) Quantification of Congo-red fluorescence intensity inside the worm body for different fibrillar forms (Number of worms 10-12).

### AL55 FL modulates VL domain aggregation across multiple germlines

To confirm that mixed aggregation modulates VL aggregation across multiple amyloidogenic germlines, we performed aggregation kinetics assays using the patient-derived AL59 and AL70 VL domains in the presence of the AL55 FL. AL59 VL and AL70 VL belong to the κ1 and λ1 germlines, respectively (Morgan et al., 2025; Bodi et al., 2009). Each VL domain (20 µM) was incubated with AL55 FL at 1:1 and 1:2 (VL:FL) molar ratios under the same experimental conditions used for AL55 VL and AL55 FL aggregation reaction. Interestingly, we found that AL55 FL markedly reduced the aggregation of both VL proteins within the timescale of our experiment. However, at longer incubation times (> 600 min), AL70 VL showed a slight increase in ThT signal at a 1:1 molar ratio (Figure S12), but complete inhibition at the 1:2 molar ratio of VL:FL. For AL59 VL, we found complete inhibition at both FL concentrations, suggesting that AL55 FL interacts with different VL domains in a non-specific manner and differentially modulates their aggregation rates (Figures 7A-C).

**FIGURE 7.**
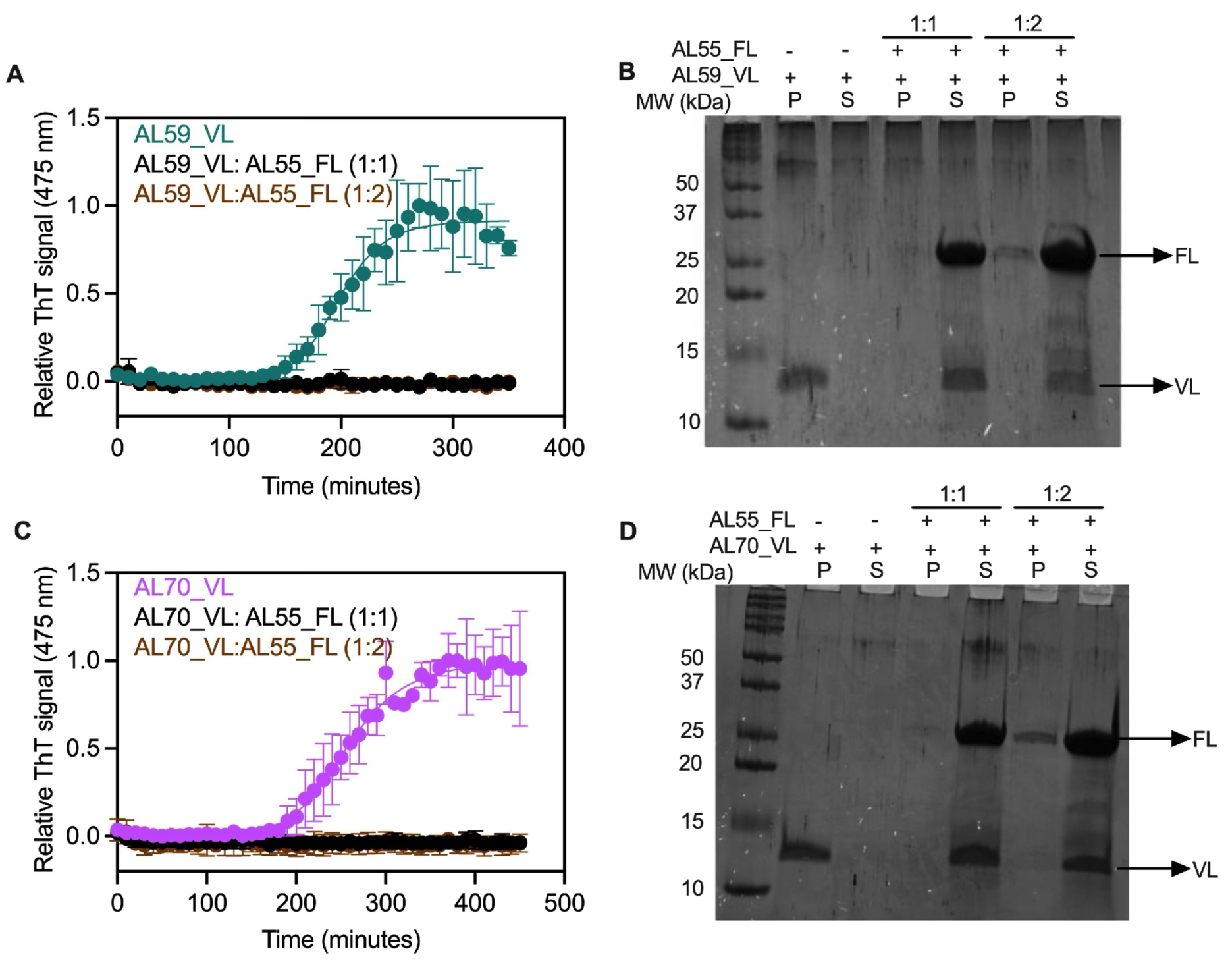
Aggregation of AL55 FL with other amyloidogenic VL domains. (A) ThT-based aggregation assays of AL59 VL (green) in the presence of AL55 FL in a molar ratio of 1:1 (black) and 1:2 (brown) (VL:FL). The experiment was performed in three technical replicates. The error bars represent the heterogeneity in ThT fluorescence emission among three replicates. (B) Supernatant and pellet fractionation of AL59 VL+AL55 FL mixed-aggregated samples. (C) ThT-based aggregation assays of AL70 VL (pink) with AL55 FL in a molar ratio of 1:1 (black) and 1:2 (brown) of VL:FL. (D) Supernatant and pellet fractionation of AL70 VL+AL55 FL aggregated samples.

These observations were further validated by supernatant–pellet fractionation of the aggregated samples after overnight incubation. In the absence of FL, both AL59 VL and AL70 VL were completely recovered in the pellet fractions, with no detectable protein in the supernatant, indicating near-complete aggregation (Figures 7B-D). In contrast, upon addition of AL55 FL, substantial amounts of both VL proteins remained in the supernatant fractions. Together, these fractionation results confirmed that AL55 FL non-specifically inhibits the aggregation of both AL59 VL and AL70 VL. This heterogeneity in the extent of modulation may arise from sequence-level differences and variation in the aggregation-competent conformational ensembles populated by individual LCs as proposed in previous studies (Meunier-Carmenate et al., 2023; Pradhan et al., 2023; Wong et al., 2024; Puri et al., 2025c). Moreover, AL55 VL does not modulate aggregation of AL70 VL, indicating the aggregation inhibition is specific to FL (Figure S13). Collectively, these findings establish that FL LCs can modulate VL aggregation across multiple amyloidogenic sequences, although the extent of modulation varies among different VLs.

## DISCUSSION

The pathological hallmark of AL amyloidosis is the extracellular deposition of structurally heterogeneous LC amyloid assemblies (Lavatelli et al., 2025; Ramirez-Alvarado et al., 2012). Proteomic analyses of ex vivo fibrillar deposits from AL patients, and from a recently developed mouse model, have consistently revealed that fibrillar extracts are a mixture of FL LCs together with their proteolytically generated fragments of varying lengths (Lavatelli et al., 2020; Lavatelli et al., 2008; Marchese et al., 2026; Martinez-Rivas et al., 2025). In contrast, high-resolution cryo-EM structures predominantly resolve ordered amyloid structures composed of VL domains, with only rare observations of partial CL-domain incorporation and no incorporation of FL LCs (Radamaker et al., 2021; Swuec et al., 2019; Radamaker et al., 2019; Radamaker et al., 2021; Puri et al., 2023; Yu et al., 2025; Yao et al., 2026; Schulte et al., 2024). Our study provides a mechanistic framework reconciling these observations by demonstrating that FL dynamically modulates VL aggregation through transient intermolecular interactions. This modulation promotes the formation of protofibrillar aggregates, which are protease-sensitive, seeding-incompetent, likely favouring their clearance and limiting their accumulation in the ECM. These species are also incompatible with high-resolution cryo-EM reconstruction.

These findings support a model in which different LC species follow divergent aggregation trajectories. Free VL domains aggregate rapidly and mature into stable, rigid, seeding-competent fibrils. The truncated CL domain interacts with neither native VL domain nor its aggregation intermediates, allowing VL to form the same fibrillar morphology as VL-alone. In contrast, FL LC engages native VL transiently and diverts it into a seeding-incompetent protofibrillar state. Non-specific sequestration of native FL by the small amount of aggregated VL might contribute to aggregation of FL in AL patients as seen in the AL mouse model where pre-formed seeds of VL domain lead to aggregation of over-expressed FL LC in heart tissues but not with only over-expressed FL protein in isolation (Matinez-Rivas et al., 2025). Moreover, the rigid VL fibrils are persistent, seeding-competent, and readily recruit soluble VL species, they form the dominant tissue deposits. These results are consistent with other aggregation studies, in which truncation, post-translational modifications and co-aggregation of Aβ-40, Aβ-42 and Tau modulate their aggregation pathways, fibrillar composition, morphology, and stability, with significant clinical consequences (Wu et al., 2022; Ozcelik et al., 2016; Mosconi et al., 2026; Puri et al., 2026). The model also offers a physiological explanation for why ex vivo cryo-EM studies predominantly resolve VL-domain-containing amyloid assemblies, while non-specifically sequestered native FL, CL, and unstable protofibrillar aggregates are excluded during sample purification for structural analysis.

MD simulation data identified the structural basis of this divergence: the stabilization of aggregation-prone regions spanning residues 38-56 and 90-103 of VL due to transient interaction with FL (Puri et al., 2025b and Puri et al., 2025c). Moreover, transient occupation of these sites by FL likely masks the primary nucleation hotspot, thereby enhancing lag time, reducing aggregation rates, promoting the formation of distinct nuclei, and consequently protofibrillar assemblies rather than canonical fibrils. The extent of this remodelling depends on the sequence and structural heterogeneity in FL LCs and truncated VL domains (Paissoni and Puri et al., 2025).

The pathological relevance of structurally distinct amyloid assemblies formed under different aggregation conditions is supported by functional toxicity studies in *C. elegans*. Rigid fibrils formed under VL and VL+CL aggregation conditions cause pronounced impairment of pharyngeal pumping. In contrast, FL-modulated protofibrillar assemblies exhibited no impairment in pharyngeal pumping. The toxicity of the fibrillar forms are significantly different than the native proteins. Among native proteins, FL is the most toxic followed by the VL and CL is not toxic. The differences in toxicity induced by native proteins and fibrillar form might be due to their different molecular mechanisms driving toxicity (Puri et al., 2025b). Together, these findings establish that LC aggregation generates both transient and persistent fibrillar populations with distinct structural and pathological properties.

Collectively, our study work supports a model in which partial proteolytic truncation of FL LC in the ECM is a prerequisite to initiate the aggregation and presence of FL and its truncated VL domain in a confined ECM space reshapes the aggregation process, fibril composition, fibril architecture, stability, and toxicity landscapes in AL amyloidosis. While heterogeneous FL-containing intermediates form during early aggregation, the stability, seeding competence, and persistence of VL-core fibrils favour their progressive enrichment within patient deposits. These findings provide a mechanistic basis for the apparent discrepancy between biochemical analyses of ex vivo aggregates and the highly homogeneous VL-core structures resolved by cryo-EM. Moreover, the concentration-dependent inhibition of aggregation of multiple VL domains by FL points towards a promising avenue for future work to use FL-derived peptide fragments as inhibitors of VL aggregation.

## MATERIALS AND METHODS

### Protein over-expression and purification

The gene encoding FL AL55, its VL and CL domains cloned in the pET21b(+) vector, was a generous gift from Prof. Stefano Ricagno, University of Milan, Italy (Puri et al., 2025b). The plasmids were transformed into the *E. coli* Rosetta (DE3) strain for protein over-expression. The overnight-grown primary culture was used for the secondary culture. Protein expression induction was performed with a final isopropyl-β-D-thiogalactopyranoside (IPTG) concentration of 0.5 mM when the optical density at 600 nm (O.D._600 nm)_ of the culture reached 0.8. The induced culture was grown for 4 h before harvesting the cell pellet by centrifugation at 6000 rpm for 20 min at 4 ℃ (Avanti-J26XP centrifuge with a JLA 8.1 rotor). Harvested cells were then used for the purification of three proteins. All proteins were purified using a common purification protocol. For purification, harvested cells were first resuspended in lysis buffer containing 20 mM Tris, 1 mM EDTA, 1 mM PMSF, and 1x Triton (pH 8.0) and lysed using sonication with an amplitude of 60% and 5 sec ON/OFF cycles for 10 min. The lysed cells were centrifuged at 12,000 rpm for 30 min at 4 ℃ using a high-speed centrifuge (Thermofisher Scientific). The inclusion bodies were collected and washed again with a buffer containing 20 mM Tris, 1 mM EDTA, and 1 mM PMSF (pH 8.0). The washed inclusion bodies were unfolded in an unfolding buffer containing 20 mM Tris, 1 mM EDTA, 1 mM PMSF, and 6 M guanidine hydrochloride (GdnHCl) for 3 h under continuous shaking. The unfolded mixture was centrifuged, and the supernatant containing the unfolded protein was then collected, and the pellet fraction containing cell debris was discarded. The collected supernatant was then mixed with refolding buffer containing 20 mM Tris, 1 mM EDTA, 1 mM PMSF, 5 mM reduced and 0.5 mM oxidized glutathione in a 1:10 v/v ratio and incubated overnight under mild shaking. The refolded samples were centrifuged to remove the aggregated protein, and the supernatant containing folded protein was filtered through 0.22 µm filters before loading onto the anion exchange chromatography (QFF 16/60, Cytiva) for further purification. The bound protein was eluted using a continuous gradient of 1.0 M NaCl. The peak fractions containing protein of interest were collected and concentrated through 3 and 10 kDa cutoff protein concentrators, depending on the size of proteins (FL AL55 dimer: app. 45 kDa, and monomeric VL and CL are 11-12 kDa each). The concentrated protein was injected into a size-exclusion chromatography (Superdex 75 10/300, Cytiva) column for final purification.

The obtained peaks of protein of interest were pooled, and the final protein concentration was measured by absorbance at 280 nm. The correct folding and oligomeric states of purified proteins were verified by collecting SEC elution volumes and circular dichroism spectra of all proteins with a typical immunoglobulin fold signature (Figure S1) (Puri et al., 2025b). The purified proteins were stored in small fractions of 500 µl at -20 ℃ for further use.

### Aggregation kinetics assay

Previous aggregation studies on the isolated AL55 VL domain from our group have established that 20 µM is an optimum concentration for its aggregation studies, and hence the same concentration was used for this work (Puri et al., 2025c). For aggregation assays in presence of FL and CL, we used 1:1 and 1:2 molar ratios of VL:FL and VL:CL proteins. Therefore, the concentrations of FL and CL were 20 and 40 µM. The aggregation conditions include 1x phosphate buffer saline (PBS, pH 7.4), 480 rpm shaking in the presence of 6 mm PTFE beads, and 10 µM Thio-flavin T dye. The temperature used for aggregation kinetics experiments was 37 ℃. Each reaction was performed in two biological and three technical replicates. The obtained data were normalized and fitted to a sigmoidal aggregation model (Equation 1), and obtained rate constants and lag times were plotted with respect to the concentration of FL and CL proteins.

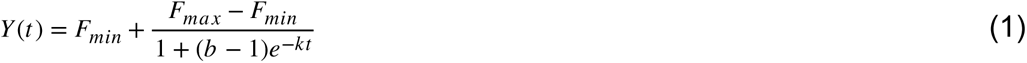

Where Y(t) is observed fluorescence intensity at time t, F_min_ is the fluorescence signal at time zero, F_max_ is the final fluorescence signal, b is a shape parameter influencing position and steepness of transition, and k is the apparent rate constant (Gade Malmos et al., 2017).

We also performed the aggregation assays of AL59 VL and AL70 VL domain proteins with AL55 FL protein. The protein concentration, aggregation assay conditions, data collection, and analysis parameters are the same as explained above. All the assays were performed in three biological replicates where each biological replicate had three technical replicates.

### Supernatant and pellet fractionation assays

To check the presence of protein in the supernatant and pellet fractions upon aggregation under different conditions, we pooled all three technical replicates in one tube and centrifuged the samples at 10,000 rpm for 30 min at 4 ℃ (Eppendorf centrifuge). The obtained supernatants were collected in a new microfuge tube, and the remaining pellets were resuspended in the same volume of buffer. 20 µl of each sample was collected and mixed with 5 µl of 5x SDS-PAGE loading dye. The samples were heated at 95 ℃ for 10 mins before loading on 12% and sometimes 15% SDS-PAGE gels. The gels were then stained with 1% Coomassie brilliant blue stain for 20 mins and destained till the clear bands appeared. The presence of different proteins in the supernatant and pellet fractions was confirmed by visual inspection of gels. This experiment was performed multiple independent times and a representative gel picture is shown in Figure 1.

To know the composition of formed fibrils after 15 days of aggregation, we set up a 300 µl aggregation reaction in an incubator shaker set at 37 ℃ and a shaking speed of 180 rpm. The different samples were VL-alone (50 µM), VL:FL (1:2), and VL:CL (1:2) prepared in 1x PBS buffer (pH 7.4). All the experiments were performed in three technical replicates. After 15 days, supernatant and pellet fractionation were done, and 20 µl of each sample was loaded on the 12% SDS-PAGE to visualise the proteins.

## TEM

TEM was used to visualize the fibrils formed under different VL alone, VL+FL, and VL+CL aggregation conditions. The samples used for TEM imaging are pellet fractions obtained from VL only, VL+FL (1:2), and VL+CL (1:2) mixed-aggregation reactions of 24 h and 15 days. We have not performed TEM imaging on FL alone and CL alone samples as both the proteins were in supernantant fraction throughout the aggregation (Figure S2). For TEM grids preparation for fibrils, the carbon-coated copper grids with 300-400 mesh size were glow-discharged using 25 mA glow discharge power for 10 sec (PELCO easiGlow). The glow-discharged grids were then incubated with 10 µl of each sample for 1 min. The residual sample and buffer were blotted with blotting paper. We then washed the grids with 10 µl of filtered Milli-Q for 30 sec before blotting. The reason for the Milli-Q wash is to remove residual PBS buffer, as it generally interferes with the uranyl acetate-based negative staining.

The washed grids were mounted with 5 µl of 0.5% uranyl acetate stain for 30 sec. The stain was then blotted, and grids were dried in a desiccator overnight before TEM data collection. The TEM data were collected at a magnification of 46 k. The acquired images were used to compare fibrillar morphologies formed under different experimental conditions.

### Cryo-EM

Cryo-EM was performed to extract high-resolution information of the protofibrils formed under VL+FL aggregation conditions. To prepare cryo-EM grids, 4 µl of a sample containing protofibrils was applied onto a glow-discharged carbon-coated copper (1.2/1.3) 300-mesh grid. The sample was blotted immediately and rapidly plunge-frozen into liquid ethane using a Vitrobot Mark IV (Thermofisher Scientific). The vitrified grids were then transferred to a Glacios microscope operating at 200 kV and equipped with an EF Falcon4i detector. A total of 7246 movies were acquired by applying an electron dose of 50 e-/Å2, which was divided into 50 frames at a magnification of 120k, corresponding to a pixel size of 0.889Å/pixel and a defocus range of -0.5 to -2.5 µm.

### Helical reconstruction

A total of 7246 multi-frame movies were dose-weighted, motion-and CTF-corrected by MotionCor2 (Zheng et al., 2017) and CTFFIND4.1 (Rohou et al., 2015), built into RELION 5.0.1 (Burt et al., 2024; He and Scheres, 2017). A total of 19969 fibrillar coordinates were manually picked from 2982 selected micrographs. A first set of 32009 particles was extracted with a particle box size of 768 pixels and an inter-box distance of 10%. The tube diameter, rise, and number of asymmetric units (NSU) were set to 150 Å, 4.8 Å, and 18, respectively. A second set of 106454 smaller segments was extracted with a box size of 280 pixel and an inter-box distance of 10%. The helical tube diameter, rise, and NSU were set to 100 Å, 4.75 Å, and 6.0, respectively. An initial and following set of reference-free 2D classes were obtained with T-regularization values of 2 and 8, respectively. The estimation resolution of 2D class averages was obtained as output for each job processing and no FSC was used for its estimation as FSC works after 3D classification and refinement. The obtained large and small 2D class averages were used to estimate the width of fibrils, distances between the β-strands, and the length of the spiral, etc., for comparison with already known information of in vitro formed AL55 VL fibrils. The data analysis pipeline is shown in Figure S5.

### PK-based limited proteolysis assay

To compare the proteolysis sensitivity of different fibrillar forms formed under VL alone, VL+FL, and VL+CL mixed aggregation experiments, the pellet fractions in an approximate concentration of 1 mg/ml were digested with 1:200 and 1:500 of protein:PK for one hour at 37 ℃ in three biological replicates. The visualisation of digestion was done on an SDS-PAGE gel. To quantify the extent of digestion among different conditions, the band intensity per lane was estimated and normalised with respect to undigested sample. Estimating the concentration for the VL+FL is always difficult due to sequestration of the FL protein along with the aggregated VL and hence average of three experiments is reported for the comparison.

### Cross-seeding aggregation studies

Seeding is an important process to enhance fibrillar growth and hence their accumulation both under in vitro and in vivo conditions. Therefore, to investigate the impact of different VL fibrillar morphologies formed under different aggregation conditions, we performed cross-seeding aggregation kinetics of the AL55 VL protein. The fibrillar seeds used for this experiment were VL-alone, VL+CL, and VL+FL aggregation reactions fibrils. To prepare the seeds, 20 µM VL alone and 1:1 molar ratios of VL+FL and VL+CL aggregation end product were sonicated in a water batch based sonicator for 30 minutes at 25 ℃. The seed percentage used was 5%, and native VL concentration in all experiments was 20 µM. The aggregation conditions set up were the same as those in the above mentioned aggregation kinetics assay. All the experiments were performed in three technical and two biological replicates. The obtained fluorescence data were normalized, plotted with respect to time, and fitted into Equation 1. The obtained rate constants of overall reactions were compared to establish the effect of seeding on aggregation rates of the VL domain.

### MD simulations

#### System preparation

Two systems were prepared for the MD simulations. We used the crystal structure (PDB: 9FVK) to prepare the system for the apo form of the VL domain (Puri et al., 2025a). We used AlphaFold3 to predict the complex of the FL dimer and the VL domain (Abramson et al., 2024). To predict the complex, we used two copies of the FL sequence and one copy of the VL sequence as input. The prediction was template-independent, with the maximum number of recycles set to 5 and the maximum number of relaxation iterations set to 200. Five structures were predicted, and the model with the lowest predicted aligned error (PAE) score was selected as the highest-confidence model.

### Molecular dynamics simulations

We performed MD simulations of the two systems: the VL domain only (VL) and the complex of VL and FL dimer (VL+FL). All simulations were performed using GROMACS 2025.4 (Abraham et al., 2015) with the Amber19SB force field (Tian et al., 2020) and the TIP3P water model (Jorgensen et al., 1983). Each system was solvated and neutralized with 150 mM NaCl to mimic physiological conditions. The systems were energy minimized using the steepest descent algorithm, followed by equilibration consisting of 125 ps in the NVT ensemble and 100 ns in the NPT ensemble, divided into two steps of 50 ns each, with position restraints on backbone atoms of 1000 kJ/mol and 500 kJ/mol. Neighbor searching was performed every 20 steps. The PME algorithm was used for electrostatic interactions with a cut-off of 0.9 nm. A single cut-off of 0.9 nm was used for Van der Waals interactions. Temperature coupling was performed with the V-rescale algorithm at 300 K. Pressure coupling was performed with the C-rescale algorithm at 1 bar (Darden et al., 1993; Bussi et al., 2007; Bernetti et al., 2020). All bond lengths were constrained using the LINCS algorithm (Hess et al., 1997). The VL+FL system was initially simulated for approximately 275 ns. The trajectory was then clustered to identify the representative structure of the largest cluster. This structure was then used for additional simulation replicas of the VL+FL interaction. Each system (complex VL+FL and apo VL) was simulated in triplicate (3 independent replicas) for a total of 1000 ns per replica with a 2 fs integration time step. All trajectories were found to be stable during the simulations (Figures S7-8).

### *C. elegans-*based toxicity assays

#### Pharyngeal pumping assay

Bristol N2 strain was propagated at 20 °C on solid Nematode Growth Medium (NGM) seeded with *E. coli* OP50 (CGC) for food. *C. elegans* pharyngeal toxicity assay was performed as previously described (Maritan et al., 2020; Diomede et al., 2014; Broggini et al., 2023). Briefly, 100 worms were suspended in 100 µl of 10 mM PBS (pH 7.4) and exposed to 100 µg/ml of VL-alone, VL+FL, and VL+CL reaction fibrils for 2 h. Worms treated with 100 µl of 10 mM PBS (pH 7.4) (vehicle) and 0.1 mM H_2_O_2_ served as the negative and positive controls, respectively. After 2 h of incubation on an orbital shaker, worms were transferred onto NGM plates seeded with *E. coli* OP50. Pharyngeal pumping rates were assessed after 4, 24, and 48 h by counting the number of terminal bulb contractions over a 30 sec interval. A total of 30 worms were analysed for each condition. Four independent biological replicates were performed. Data were analysed using GraphPad Prism software by one-way ANOVA followed by Bonferroni’s *post hoc* test. A *p*-value of <0.05 was considered significant.

### Fibril uptake assays

To visualize fibril uptake, L4-stage worms were fed Congo red-stained fibrils for 2 h in 10 mM PBS (pH 7.4). Following incubation, worms were washed once with 10 mM PBS (pH 7.4), anesthetized in M9 buffer containing 50 mM sodium azide, and mounted on 2% agarose pads. Images were acquired using an SMZ18 stereo fluorescence microscope, with 10–12 worms imaged for each condition. For fluorescence quantification, individual worms were manually outlined in ImageJ, and the fluorescence intensity, area, integrated density, and mean background fluorescence were measured. Data were analyzed using GraphPad Prism software with an unpaired two-tailed t-test; p-value < 0.05 (*) was considered significant, and ns denotes a non-significant difference.

## SUPPLEMENTARY MATERIAL DESCRIPTION

**Figure S1**: Model proteins; **Figure S2**: Pellet and Supernatant fractionation of CL and FL protein after overnight aggregation. **Figure S3**: Quality check of the purified FL, VL, and CL proteins; **Figure S4**: Aggregation kinetics assays raw data; **Figure S5**: Aggregation kinetics assay of VL in presence of BSA. **Figure S6**: Representative TEM images after 24 h of aggregation reaction; **Figure S7**: CryoEM data analysis pipeline; **Figure S8**: Composition of co-aggregated fibrils; **Figure S9:** SDS-PAGE showing the sensitivity of VL alone, VL+FL, and VL+CL fibrils against proteinase K digestion. **Figure S10**: Cartoon representation of the modeled VL+FL complex, with the VL-VL interaction region highlighted and enlarged; **Figure S11**: Backbone RMSD plots; **Figure S12**: Aggregation kinetics raw data of AL59 and AL70 VL domains in presence of AL55 FL. **Figure S13**: Aggregation kinetics of AL70 VL domain in presence of AL55 VL and AL55 VL at 20 and 100 μM protein concentrations.

## AUTHORS CONTRIBUTIONS

**Basudha Patel and Sharvari Palkar:** Purified proteins, conducted the aggregation assays, cross-seeding aggregation assays, proteolysis assays and TEM data collection. **Shivansh Singh, Twinkle Aggarwal, Swasti Pradhan, Saara Syam, Sandhya Gokula Nathan**: Cryo-EM data analysis, aggregation assays, proteolysis and SDS-PAGE analysis. **Amit Kumawat**: Molecular dynamics simulations, funding acquisition, and writing and editing the manuscript. **Annesha Ghosh**: performed *C. elegans*-based toxicity assays and wrote and edited the manuscript. **Jogender Singh**: supervised toxicity assays, funding acquisition, and editing the manuscript. **Sarita Puri:** designed the research, conducted experiments, manuscript writing and editing, funding acquisition, and supervised the project.

## Supporting information

/Users/sarita/Desktop

## ACKNOWLEDGEMENTS

This work was supported by DST-INSPIRE Faculty fellowship (DST/INSPIRE/ FACULTY-2023/IFA-23-LSBM-274) and ANRF-PMECRG grant (ANRF/ECRG/ 2025/000226/LS, Govt. of India to S.P.; A.K has been funded by e.INS-Ecosystem of Innovation for Next Generation Sardinia (cod. ECS 00000038) funded by the Italian Ministry for Research and Education (MUR) under the National Recovery and Resilience Plan (NRRP) -MISSION 4 COMPONENT 2, CUP F53C22000430001 University of Cagliari, Italy. A.K. acknowledges CINECA ISCRA for access to the LEONARDO supercomputer, owned by the EuroHPC Joint Undertaking and hosted by CINECA (Italy), under the projects Class C: HP10CWQCG8 and Class B: HP10BEOUG4. The *C. elegans* work was supported by the Har-Gobind Khorana-Innovative Young Biotechnologist Fellowship to J.S (File No. HRD-17011/2/2023-HRD-DBT). We acknowledge Prof. Jayant Udgaonkar and his lab members for their constant support and valuable discussions throughout this study. We acknowledge Prof. Gayathri Pananghat for her valuable support and discussion on the study. We thank Mr. Kalyane Shubham Shivaji for helping us with TEM and cryo-EM data collection. We also acknowledge the Biology Department, IISER Pune, and the cryoEM facility, IISER Pune, for providing the infrastructure for our experiments. We also acknowledge Prof. Stefano Ricagno, University of Milan, Italy, for giving us plasmids encoding FL, VL, and CL proteins.

## DATA AVAILABILITY STATEMENT

The MD simulation data are available in the Zenodo repository with the following doi: 10.5281/zenodo.21677764. Rest of the data supporting this study can be found in the main text and supporting information files of the manuscript.

## AI DISCLOSURE STATEMENT

AI was not used while preparation of this manuscript.

