## Supplementary material for "Antibody full-length light chain reshapes variable domains aggregation without altering their dominance in amyloid fibrils": /Users/sarita/Desktop: Supplementary figures_final.pdf

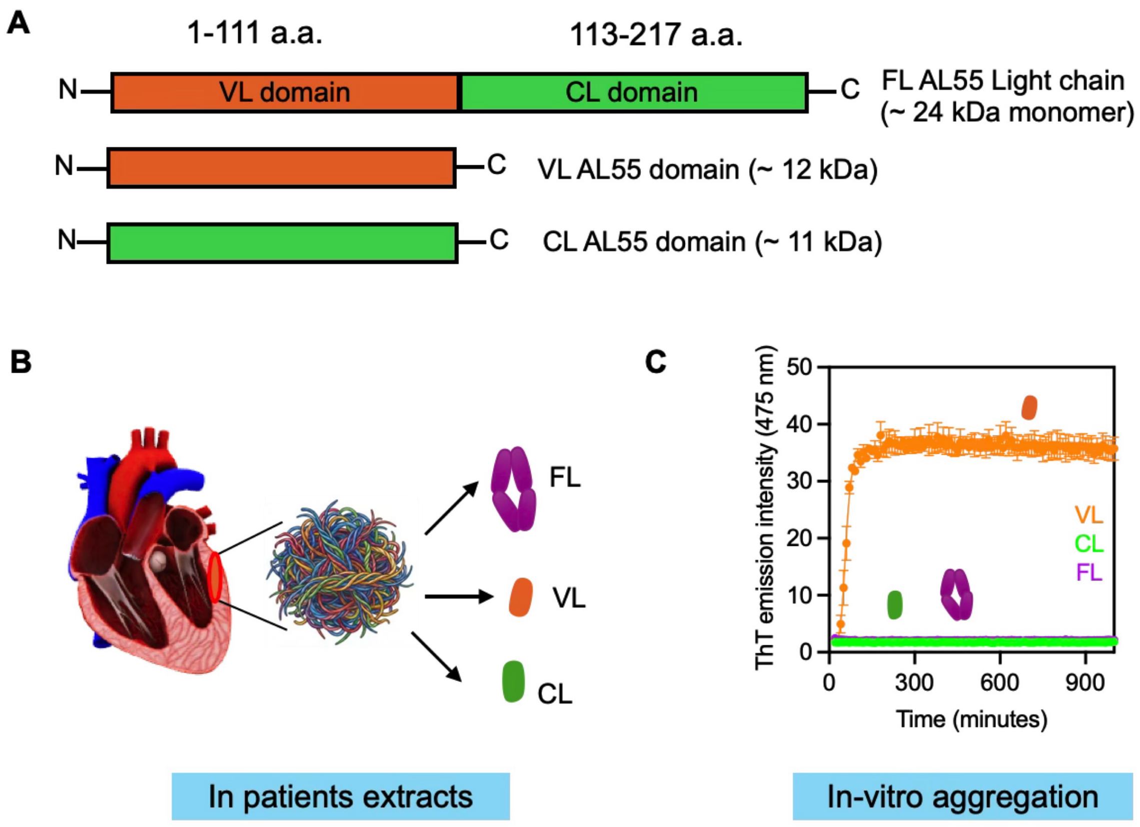


**Figure S1. Model proteins**. (A) Line diagram showing organisation of VL (orange) and CL (green) domains in FL AL55 protein. (B) Pictorial representation of mixed aggregates containing FL, VL, and CL proteins found in AL patients. (C) In vitro aggregation profiles of recombinantly purified FL AL55 (magenta), VL (orange), and CL (green) proteins.

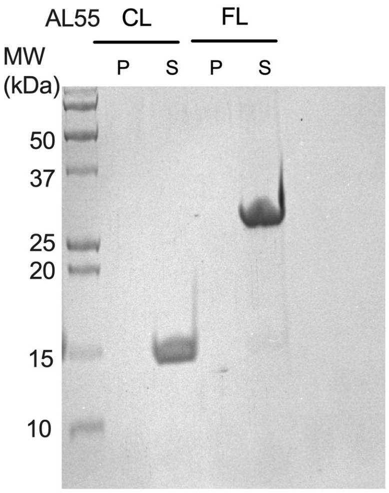


**Figure S2. Supernatant and pellet fractions of CL and FL protein after 24 h aggregation reaction.** As these proteins do not aggregates in our aggregation conditions, 100% of both proteins were present in the supernatant fraction.

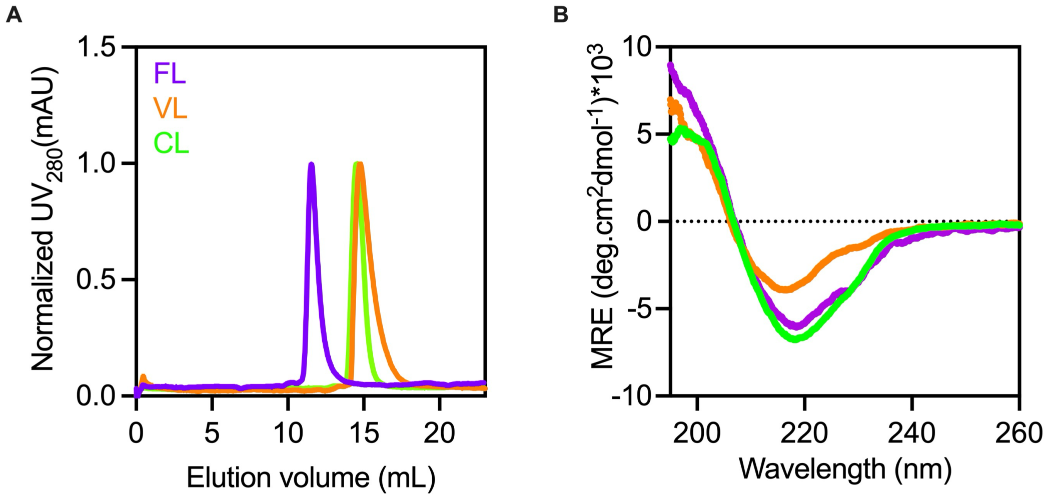


**Figure S3. Quality check of the purified FL, VL, and CL proteins**. (A) Size-exclusion chromatography profiles of purified FL (magenta), VL (orange), and CL proteins (green). (B) Far-UV CD spectra of FL, VL, and CL proteins.

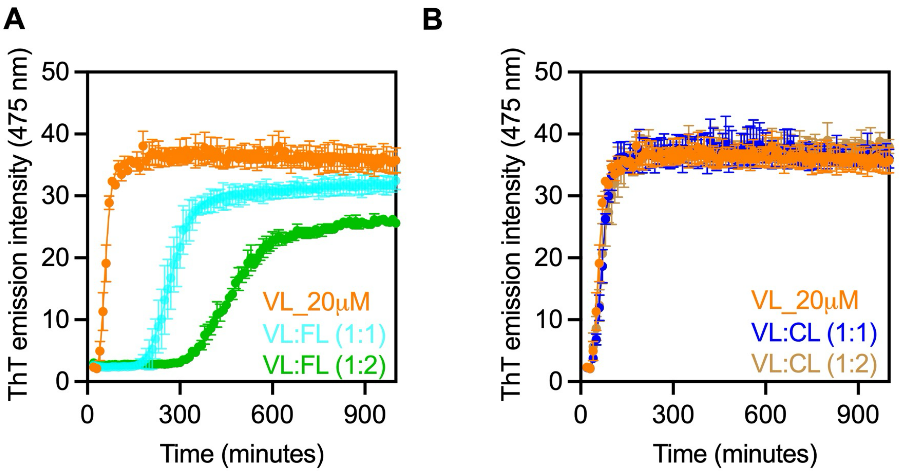


**Figure S4. Aggregation kinetics assays raw data**. (A) Aggregation kinetics raw data of VL: FL protein at a molar ratio of 1: 1 and 1: 2 (VL: FL). (B) Aggregation kinetics raw data of VL: CL protein at a molar ratio of 1: 1 and 1: 2 (VL: CL).

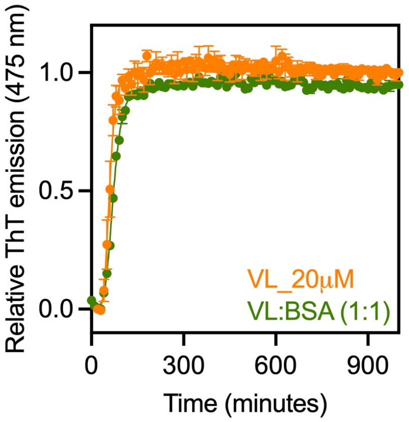


**Figure S5. Aggregation kinetics assay** of VL alone (orange) and in presence of BSA (1:1 molar ratio) (green). The protein concentration of each protein was 20 μM. All experiments were performed in three technical replicates.

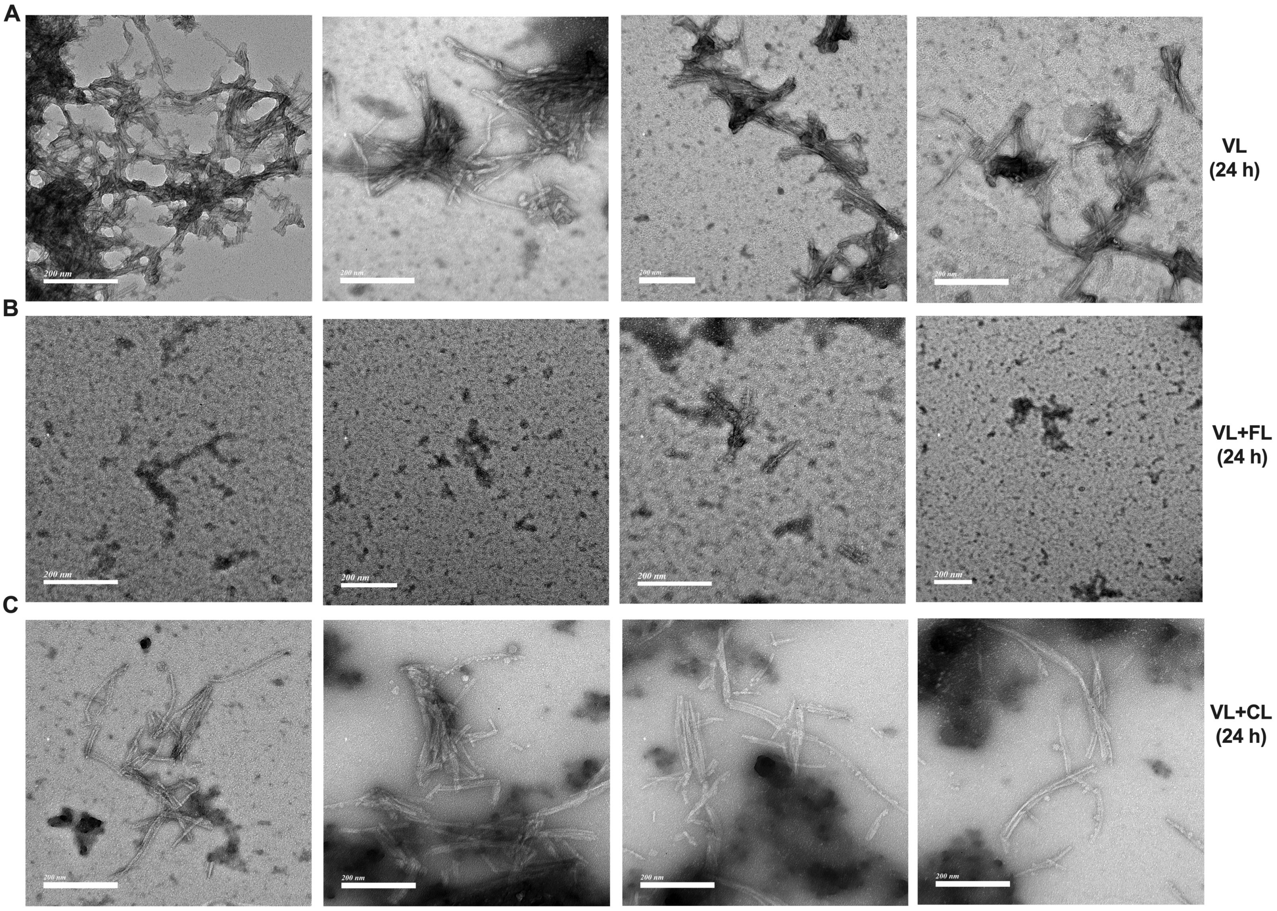


**Figure S6. Representative TEM images after 24 h of aggregation reaction**. (A) TEM images of VL aggregates. (B) VL+FL co-aggregated sample. (C) VL+CL co-aggregated sample. We don’t see any fibrils or amorphous aggregates in the FL and CL alone samples and hence their micrographs are not included here.

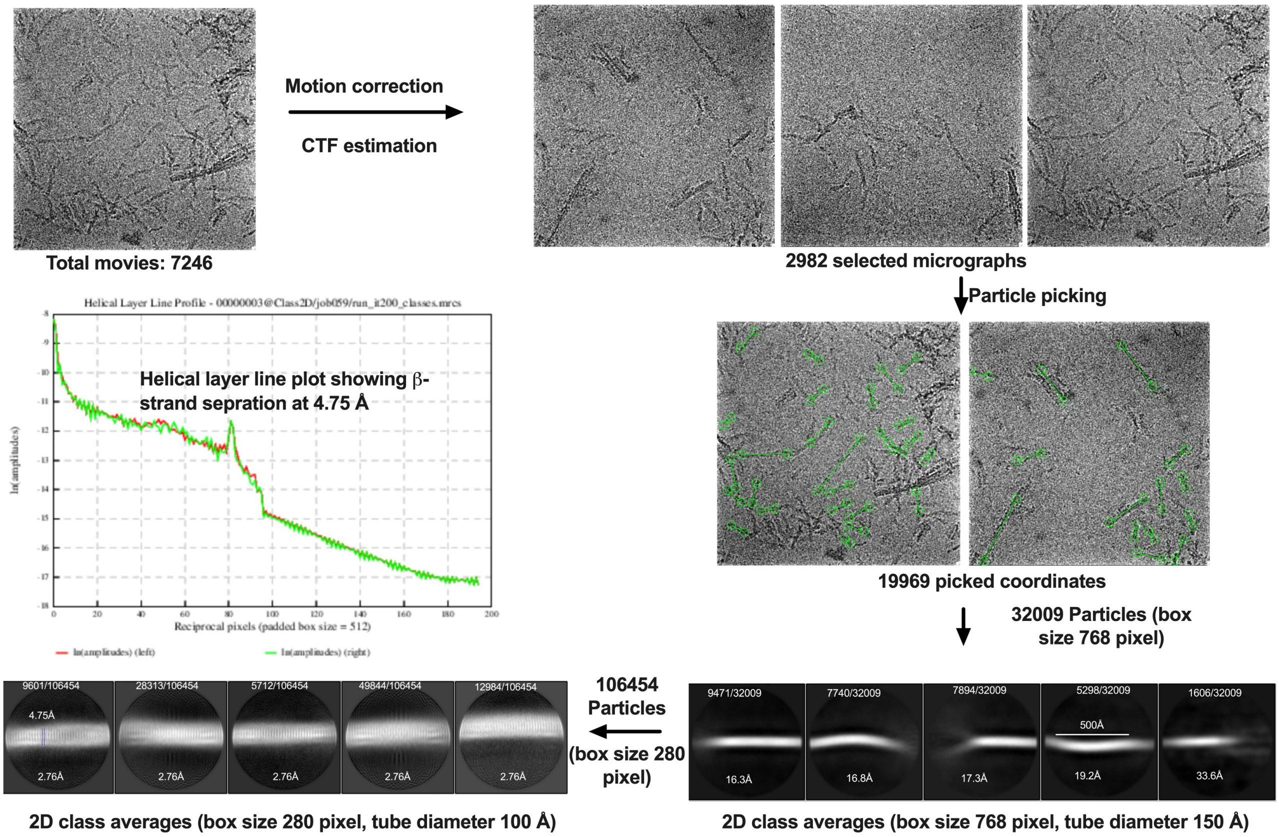


**Figure S7. CryoEM data analysis pipeline** showing motion corrected and CTF estimated micrographs, coordinate picking, particle extractions with box size of 768 and 280 pixels. Large 2D class averages to get the helical parameter information. Small 2D class averages to get high resolution information of individual β-strand separation. Helical layer line plot showing a typical β-strand separation of 4.75 Å.

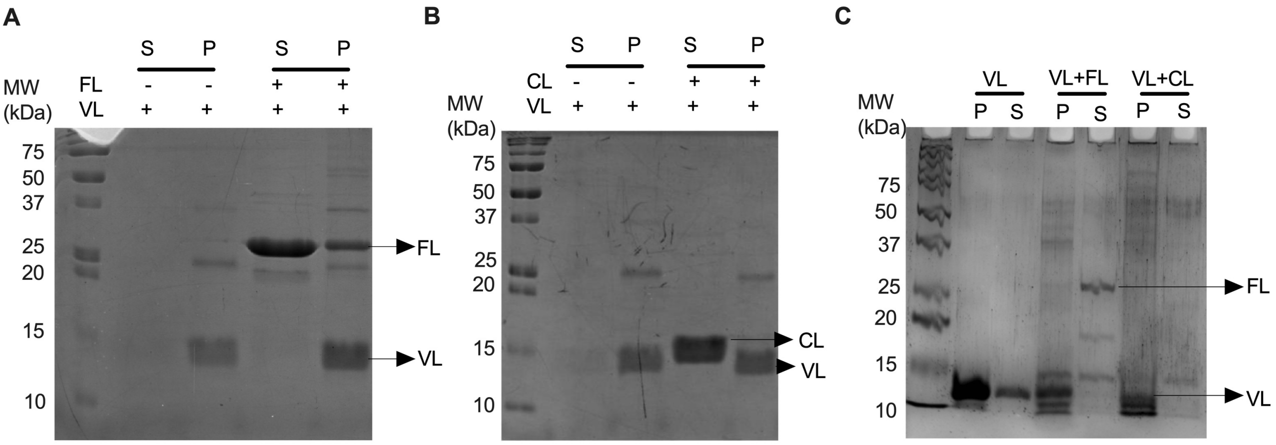


**Figure S8. Composition of co-aggregated fibrils**. (A) Supernatant and pellet fractionation of VL alone, VL+FL co-aggregated sample after 24 h of aggregation. (B) Supernatant and pellet fractionation of VL alone, VL+CL co-aggregated sample after 24 h of aggregation (C) Supernatant and pellet fractionation of VL alone, VL+FL, and VL+CL co-aggregation after 15 days of aggregation. The samples details is labelled on each lane on the gel.

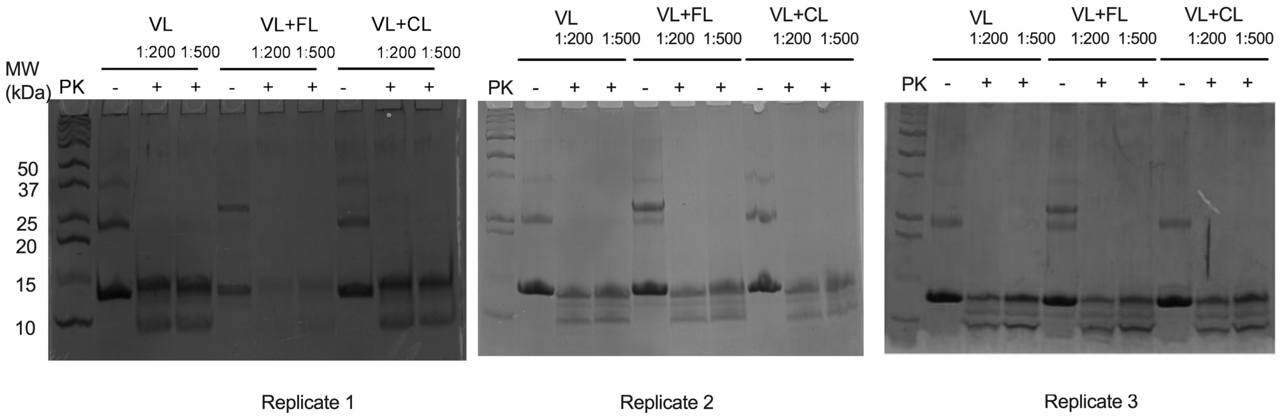


**Figure S9: SDS-PAGE showing the sensitivity of VL and different mixed aggregation reaction fibrils (VL+FL, VL+CL) against proteinase K digestion**. Three SDS-PAGE are three independent proteolysis experiments carried out from three different days aggregation end products. The loading order in all gels are same which include; Lane 1 is a medium-range molecular weight marker. Lane 2 is VL fibrils without PK treatment. Lanes 3 and 4 are PK-digested VL fibrils at a ratio of 1:200 and 1:500 (PK:protein), respectively. Lane 5 is VL+FL aggregates without PK digestion. Lanes 6 and 7 are the VL+FL aggregates digested with a 1:200 and 1:500 PK:protein ratio, respectively. Lane 8 is VL+CL fibrils, and lanes 9 and 10 are the PK-digested VL+CL fibrils with 1:200 and 1:500 PK:protein ratios, respectively.

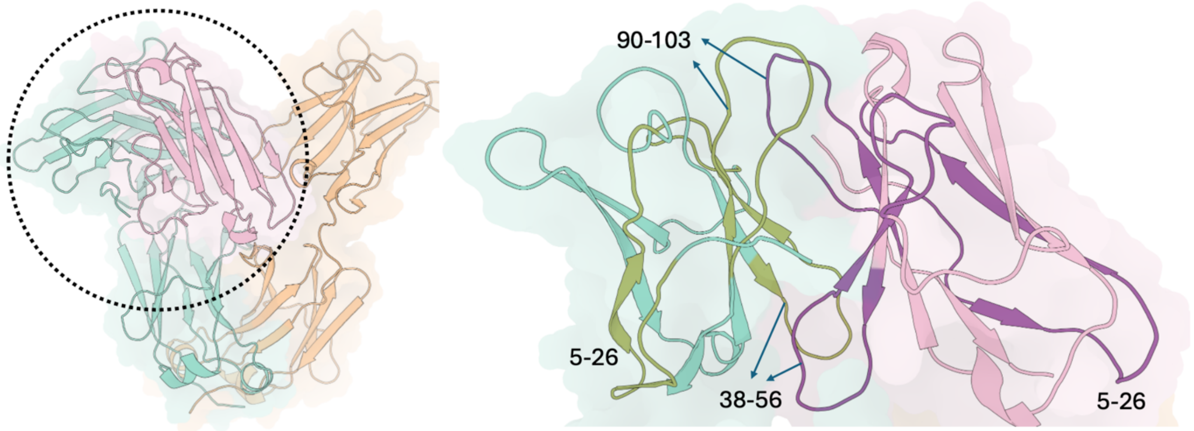


**Figure S10. Cartoon representation of the modeled VL+FL complex, with the VL-VL interaction region highlighted and enlarged.** The VL domain from the FL dimer is shown in teal, and the interacting VL domain is shown in pink. Aggregation prone regions (APRs) such as residues 5-26, 38-56, and 90-103 are highlighted in green in the FL-associated VL and in violet in the interacting VL domain.

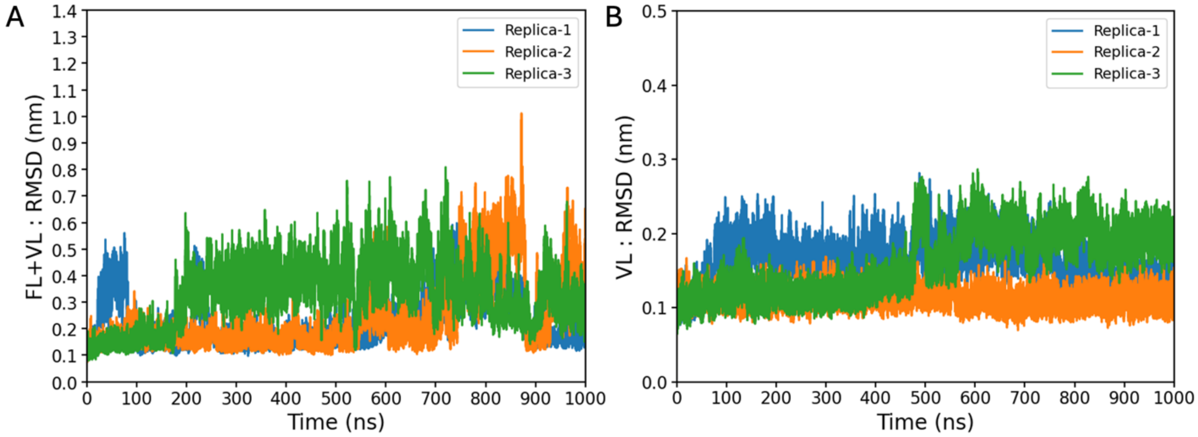


**Figure S11. Backbone RMSD plots** of the (A) VL+FL complex and (B) *apo* VL, across three independent replicas.

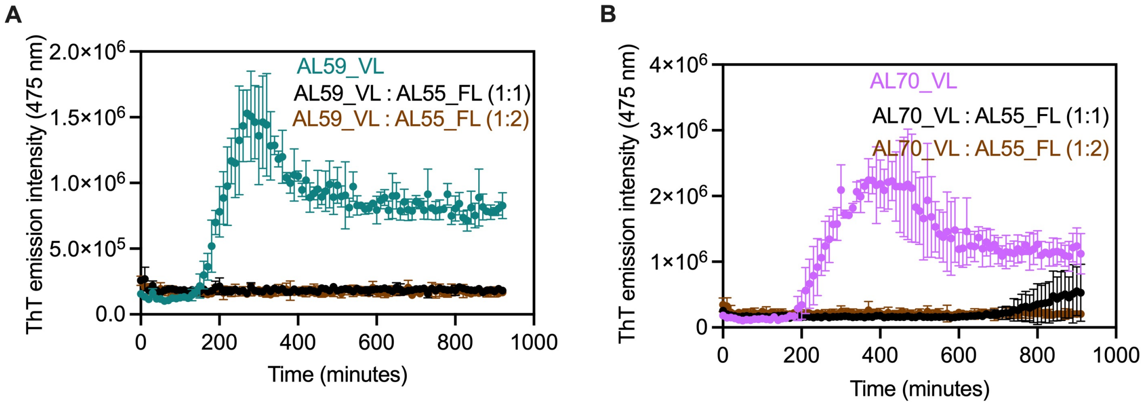


**Figure S12. Aggregation kinetics raw data of AL59 and AL70 VL domains in presence of AL55 FL**. (A) Raw data showing aggregation kinetics of AL59_VL in presence of 1:1 and 1:2 molar ratio of AL59_VL: AL55_FL. (B) Raw data showing aggregation kinetics of AL70_VL in presence of 1: 1 and 1: 2 molar ratio of AL70_VL: AL55_FL.

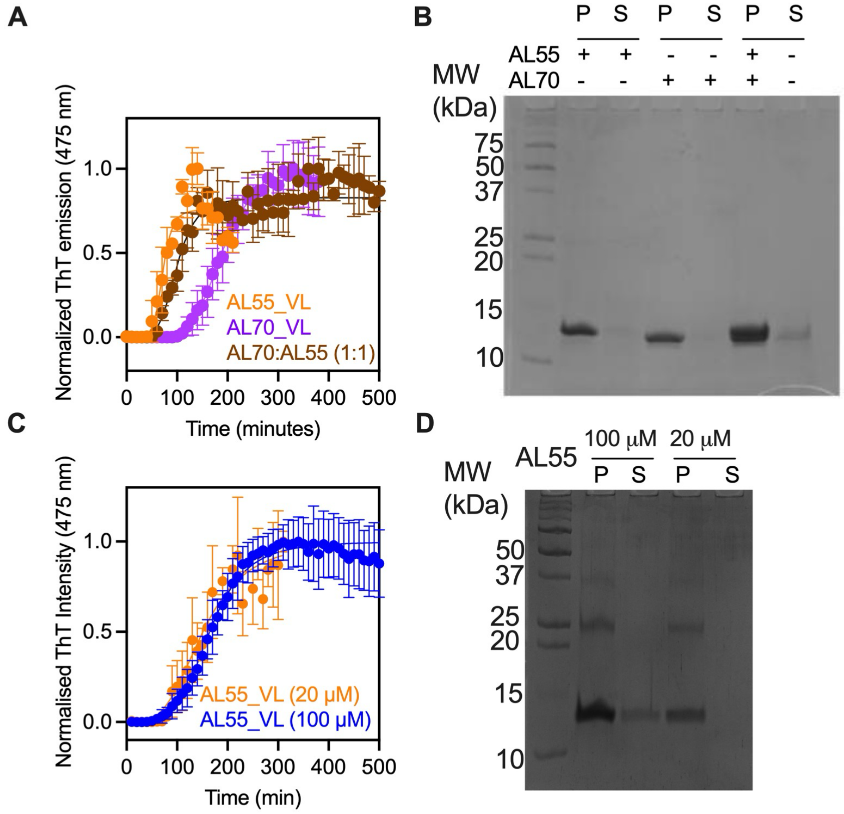


**Figure S13. Aggregation kinetics of AL70_VL domains in presence of AL55_VL and AL55 VL at different concentrations**. (A) Aggregation kinetics of AL55_VL, AL70_VL and a mixture of AL55:AL70 VL domains in a molar ratio of 1:1. (B) SDS-PAGE showing amount of proteins present in the pellet and supernatant fractions collected at the end of aggregation reactions. (C) Aggregation kinetics of AL55 VL at 20 and 100 μM protein concentrations. (D) SDS-PAGE showing amount of protein present in the pellet and supernatant at the end of the aggregation reaction.
